# Non-parametric estimation of population size changes from the site frequency spectrum

**DOI:** 10.1101/125351

**Authors:** Berit Lindum Waltoft, Asger Hobolth

## Abstract

The variability in population size is a key quantity for understanding the evolutionary history of a species. We present a new method, CubSFS, for estimating the changes in population size of a panmictic population from the site frequency spectrum. First, we provide a straightforward proof for the expression of the expected site frequency spectrum depending only on the population size. Our derivation is based on an eigenvalue decomposition of the instantaneous coalescent rate matrix. Second, we solve the inverse problem of determining the variability in population size from an observed SFS. Our solution is based on a cubic spline for the population size. The cubic spline is determined by minimizing the weighted average of two terms, namely (i) the goodness of fit to the SFS, and (ii) a penalty term based on the smoothness of the changes. The weight is determined by cross-validation. The new method is validated on simulated demographic histories and applied on data from nine different human populations.

## Introduction

The variability in population history is informative about the evolution of a species. For the estimation of the variability in population size, The 1000 Genomes Project Consortium (2015) used a two-step estimation procedure to obtain smooth curves. First, a piecewise constant population size is estimated for one or more individuals from the populations. Second, a smoothing spline is applied to produce visually appealing curves. We avoid this two-step procedure by tailoring the theory of smoothing splines directly to the inference problem.

We propose a novel non-parametric method CubSFS for estimating the changes in population size based on the site frequency spectrum (SFS). Polanski et al. (2003a) and Polanski and Kimmel (2003b) provide a method for calculating the expected SFS given the changes in population size. We aim to solve the inverse problem: Given an observed SFS we want to estimate the changes in population size. Myers, Fefferman, and Patterson (2008) showed that the solution to this problem is in general not unique (but see Bhaskar and Song (2014)), and we therefore need to regularize or constrain the variability in population size.

Denote the SFS by *ξ* = (*ξ*_1_, *ξ*_2_, *…, ξ*_*n-*1_) such that *ξ*_*i*_ is the number of segregating sites with *i* derived alleles among the *n* sampled sequences. The population size *N*(*r*) is the number of diploid individuals in the population at generation *r*. We use scaled time *t* where one time unit corresponds to 2*N* (0) generations. In this time *λ*(*t*) = *N*(0)/[*N*(2*N*(0)*t*)] is the coalescent rate at time *t*. The integrated intensity is defined as

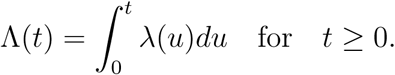

The main aim is to estimate a curve Λ = {Λ(*t*): *t* ≥ 0} that fits the data and at the same time is regular. It is straightforward to transform the integrated intensity to variability in population size.

The CubSFS method uses a roughness penalty approach (Green and Silvermann, 1994) to solve the inverse problem (see Figure 1). The main idea is to estimate the variability in population size by weighing two opposing forces: The similarity between the expected and the observed SFS versus a slowly varying curve.

**Figure 1:**
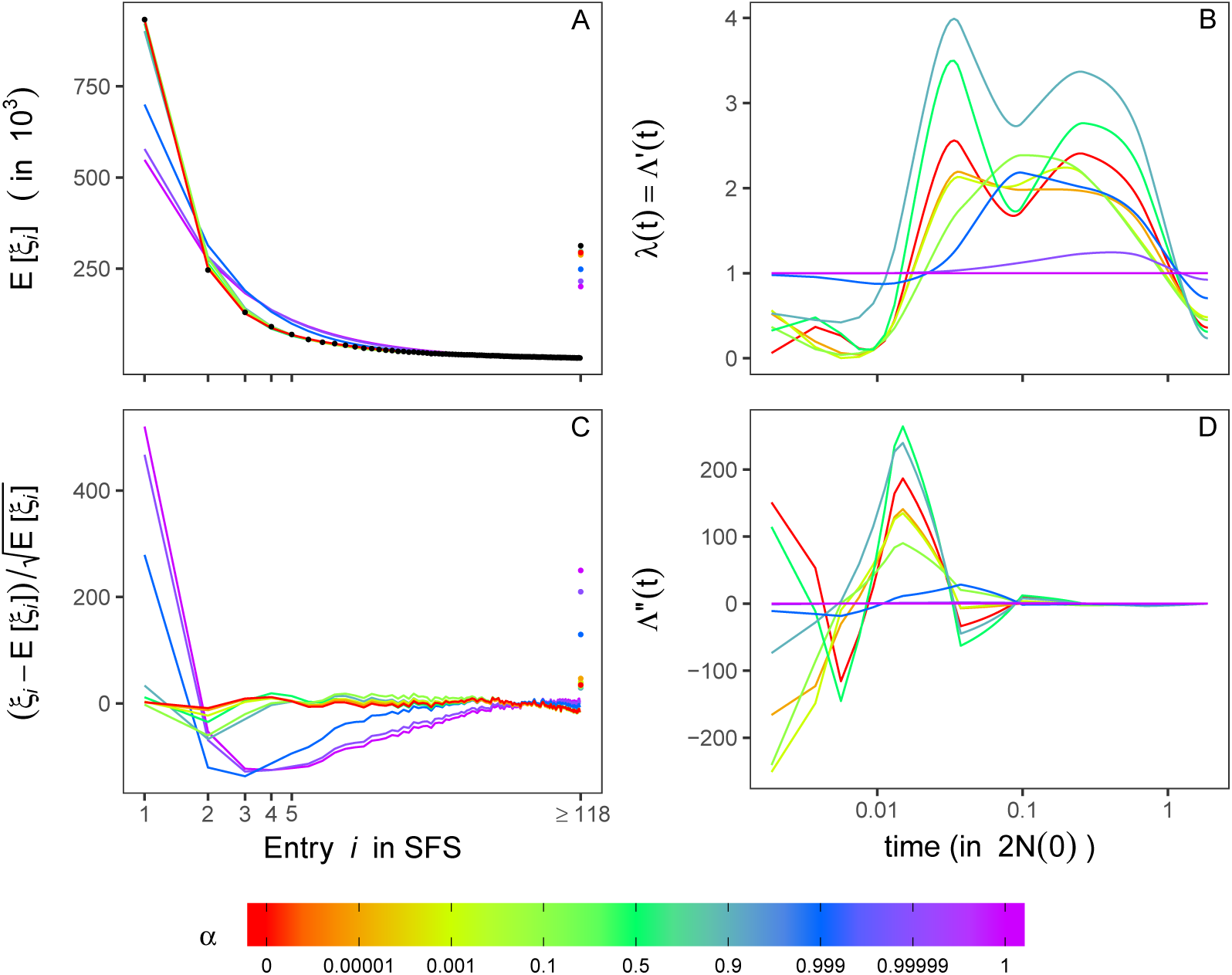
The elements of the score function. The effect of the smoothing parameter *α* on the elements of the score function (see equation (1)) for the CEU population (*m* = 14 and *t*_*m*_ = 1.875). To the left, is A: The expected and the observed SFS (black dots), and C: The residuals which is the square root of the relative distance to the observed SFS. For both plot the tail is grouped to include at most 10% of the sites. To the right, is B: The coalescent rate, and D: The derived coalescent rate with time in coalescent units. Notice that the colour legend is not linear.

In particular our solution is the function that minimizes the score function

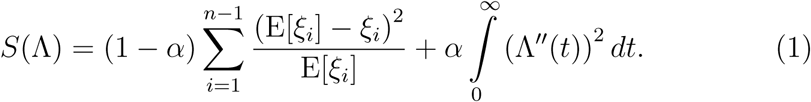

The first term in the score function is the sum of squares of the residuals 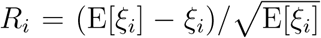 measuring the distance between the observed and the expected SFS. Here E[*ξ*_*i*_] is the expected number of sites with *i* derived alleles determined from ∧ using the theory from Polanski et al. (2003a) and Polanski and Kimmel (2003b). The second term 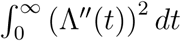 is a roughness penalty, and the smoothing parameter 0 *< α <* 1 determines the amount of regularization.

If *α* is large then the most important term in the score function is the roughness penalty and the minimizer 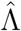 will display very little curvature. On the other hand if *α* is small then the main contribution to the score function is the residual sum of squares, and the curve estimate 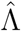must resemble the data even if it requires a rather variable curve. The smoothing parameter is estimated using cross-validation.

In Figure 1 we illustrate the method. For a range of values of *α* we show the expected site frequency spectrum (top left), the corresponding residuals *R*_*i*_ (bottom left), the coalescent rate (top right), and the roughness penalty (bottom right) for the estimated value of Λ.

The SFS is a popular summary statistics for genetic variation and in the last few years multiple methods have been developed to infer population histories from the SFS. These methods include estimates of the changes in population for a single population (Reppell et al., 2014; Eldon et al., 2015; Liu and Fu, 2015; Bhaskar et al., 2015; Gao and Keinan, 2016), and estimates of ancestral population sizes, split times and migration rates using the joint SFS for multiple populations (Gutenkunst et al., 2009, 2010; Lukic and Hey, 2011, 2012; Excoffier et al., 2013).

The SFS approach most often assumes independence between the segregating sites. Methods that take recombination or linkage into account have recently been developed. Palacios (Palacios and Minin, 2013; Palacios et al., 2015) and Lan et al. (2015) develop a method based on gene genealogies. They provide a non-parametric Bayesian estimate of the population size using a Gaussian process as a prior on the coalescent rate variability. However, the gene genealogies are not directly available and the error of inferring these have large effects on the estimated population size (Palacios et al. (2015)). Hidden Markov models (HMMs) are a popular framework for integrating out the unknown gene genealogies. The HMM framework for estimating changes in population size is well known in the PSMC method (Li and Durbin, 2011), the MSMC method (Schiffels and Durbin, 2014), The diCal method (Sheehan et al., 2013), and the SMC++ method (Terhorst et al., 2017).

Our CubSFS method shares many properties with the SMC++ method of Terhorst et al. (2017), however, here we describe some of the subtle differences. Terhorst et al. (2017) also regularize the variability in population size using a roughness penalty approach with penalty score 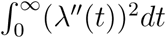. We suggest the penalty 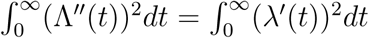, since if the weight on the penalty is high, then the score function is minimized if *λ*'(*t*) ≃ 0 for *t* ≥ 0, implying an approximately constant population size. The smoothing parameter is a central parameter for our method, and is estimated using a cross-validation procedure. This parameter is chosen from experience in the SMC++ method. Finally, in the CubSFS method we pay detailed attention to the number and type of time intervals. Especially, we are able to choose the number of time points by means of the Akaike Information Criteria.

An alternative method is based on the Approximate Bayesian Computation (ABC) approach (Boitard et al., 2016). Here, the idea is to simulate changes in population size, calculate the corresponding appropriate summary statistics, and compare with the observed data summary statistics. If the simulated data is similar to the observed data, then the population size history is accepted. Finally, the posterior distribution of the population histories is determined based on the accepted samples.

We show that CubSFS is an attractive alternative to the methods mentioned above. In particular we can handle a multitude of samples through the SFS. Further, we avoid simulation-based methodology and prior parametric assumptions by detailed analytical considerations. Our method is validated through simulations and applied to data from nine different populations from the 1000 Genomes project (The 1000 Genomes Project Consortium, 2015).

## Results

### Inference on 9 different populations

We estimate the variability in population size for 9 different populations using the CubSFS method. The results are shown in Figure 2 (additional results are shown in Supplementary Figures S16 - S24). The results are very similar within continents Asia (CHB, CHS, and JPT), Europe (CEU, FIN, GBR, and TSI) and Africa (LWK and YRI). The Asian and European populations agree on an overall bottleneck ranging from 2 to 300 thousand years ago, with a possible slightly increase in populations size approximately 20 - 30 thousand years ago. The African populations agree on one bottleneck coherent with the decline in population size of the other 7 populations happening furthest back in time. The African bottleneck is indicated slightly earlier than the other 7 populations, agreeing with the out of Africa hypothesis.

**Figure 2:**
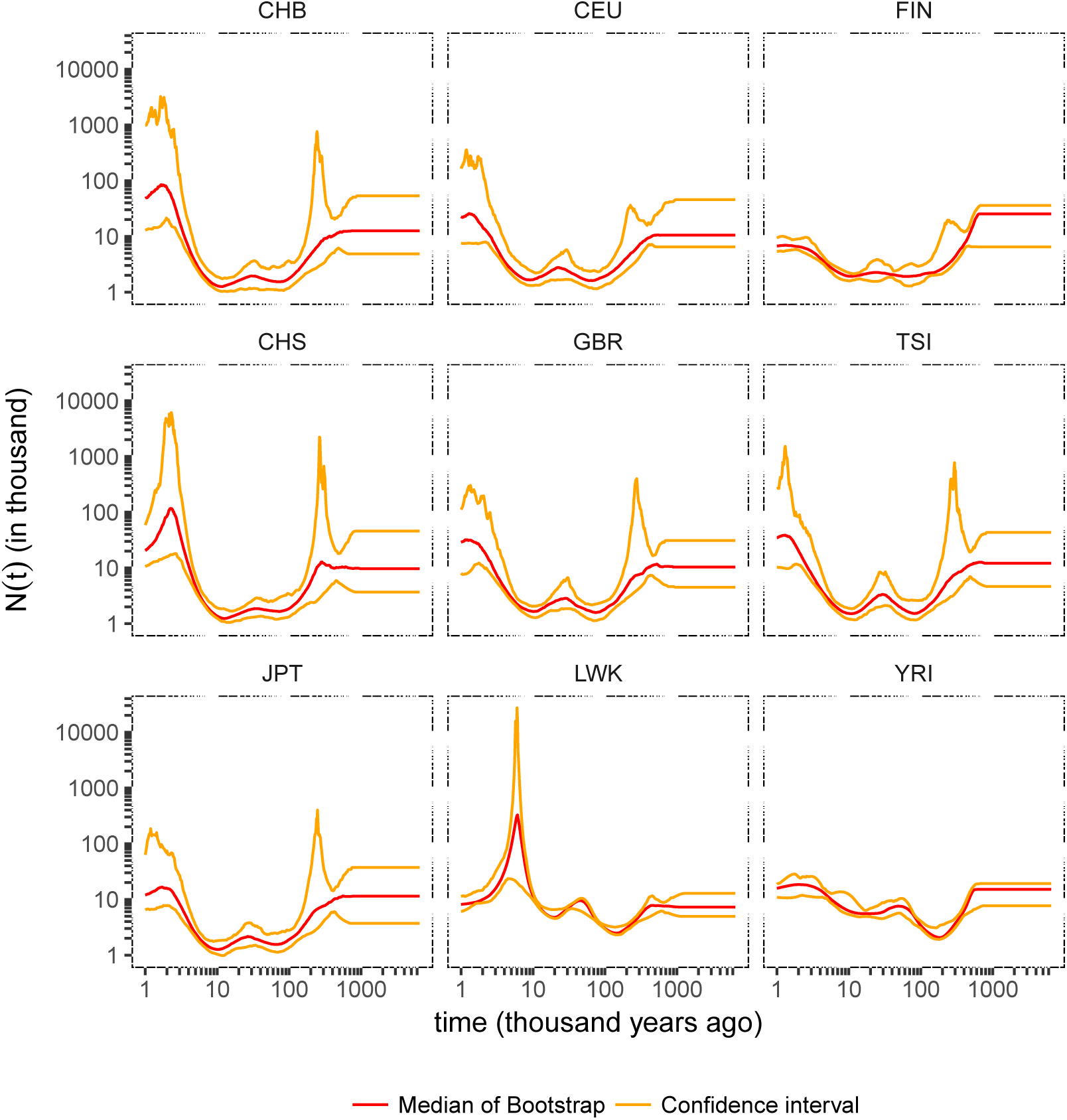
The final choice for the 9 different populations. The red line is the point-wise median of the bootstrap estimates using the CubSFS method. The orange lines are the point-wise confidence intervals. The final choice of parameter setting is determined by the AIC based on the point-wise median.

### Simulation studies

To verify our method we consider four different scenarios: Exponential growth, two epochs, a bottleneck, and a zigzag model (Schiffels and Durbin, 2014). We consider 200 sequences of length 5 · 10^8^ and the number of segregating sites is 6 · 10^6^.

The population size inferred from CubSFS is shown in Figure 3 (additional results are provided in Supplementary Figures S6 - S10). The method generally provides sensible results for all models. The exponential growth model is inferred particularly well. For instantaneous changes as the two epochs, the bottleneck, and the zigzag models the smoothing seems to detect the trends in general, but the instantaneous changes in population size are more difficult to detect.

**Figure 3:**
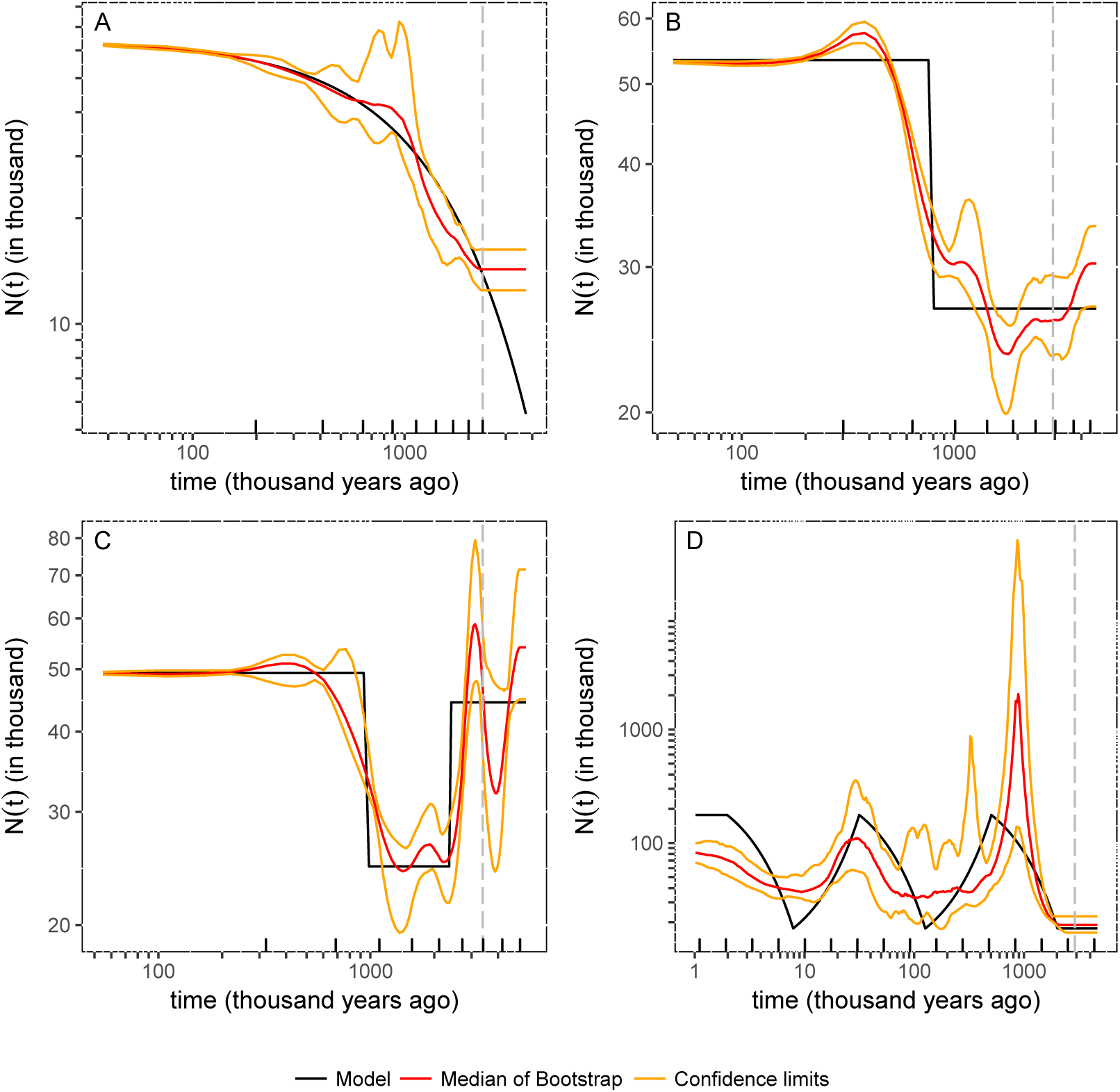
The final choice for the simulated models. The black line is the true model and the red line is the point-wise median of the bootstrap estimates based on the CubSFS method. The orange lines are the point-wise confidence intervals. The dotted grey vertical lines are the expected time to most recent common ancestor for each of the four models. The rug points are the time points used by the CubSFS method. For three of the models A: exponential growth, B: two epochs, and C: bottleneck the times points are placed linearly on a logarithmic scale in coalescent units prior to estimation. For model D: zigzag the time points are placed according to knowledge of the model. The final choice of parameter setting is determined by the AIC based on the point-wise median of the results from the bootstrap samples.

## Methods

### Theory

Recall that *ξ* = (*ξ*_1_, *ξ*_2_, *…, ξ*_*n-*1_) is the SFS for *n* sequences. Furthermore *N*(.) is the population size, *λ*(*t*) = *N*(0)*/N*(2*N*(0)*t*) is the coalescent rate, and 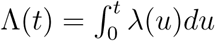 is the integrated intensity. The score function is given by

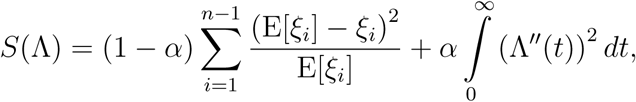

where 0 *< α <* 1 is the smoothness parameter, and E[*ξ*_*i*_] is the expected number of sites with *i* derived alleles based on the integrated intensity function Λ.

The value E[*ξ*_*i*_] is evaluated using the probability *p*_*i*_(Λ) of observing a site with *i* derived alleles as

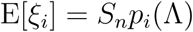

where *S*_*n*_ is the number of segregating sites,

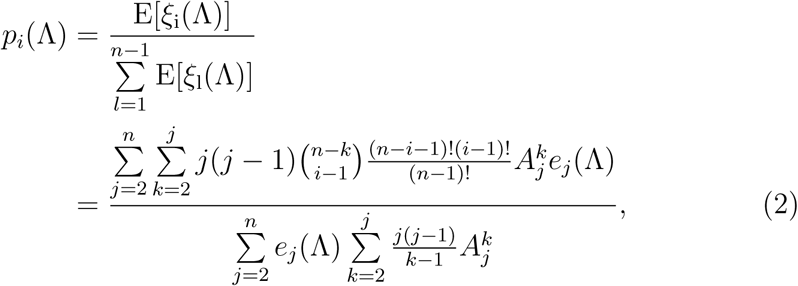

and 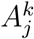 and *e*_*j*_(Λ) are defined by Polanski et al. (2003a):

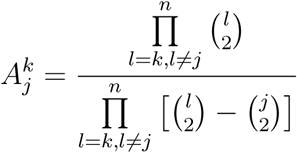

for *k ≤ j ≤ n*, 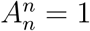, and

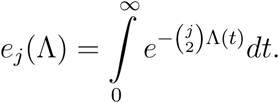

Equation (2) can be evaluated using constants defined by Polanski and Kimmel (2003b). A straightforward proof of equation (2), different from that provided by Polanski et al. (2003a), is given in the Supplementary Information.

We estimate the integrated intensity by a cubic spline (Green and Silvermann, 1994) defined on *m* + 1 time points back in time (see Supplementary Figure S2). The time points 0 = *t*_0_ ≤ *t*_1_ ≤ … ≤ *t*_*m*_ are proposed to be evenly distributed on a logarithmic scale (Li and Durbin, 2011), however, the placement should incorporate any prior knowledge of the model. The cubic spline has the following properties (see also the Supplementary Information):

i) *λ*(0) = Λ′(0) = 1
ii) Λ (0) = *a*_0_ = 0
iii) after *t*_*m*_ the intensity is constant
iv) the integrated intensity is increasing.

### Implementation

Given *α*, *t*_0_, *t*_1_, *…, t*_*m*_ and the values Λ(*t*_*i*_) = *a*_*i*_ for *i* = 0, *…, m* the cubic spline of Λ is fully specified (see the Supplementary Information). The score function (1) is minimized with respect to the values of 0 = *a*_0_ ≤ *a*_1_ … ≤ *a*_*m*_ given *α* and *t*_0_, *t*_1_, *…, t*_*m*_. The search in (*a*_1_ ≤ … ≤ *a*_*m*_), determining an increasing cubic spline that minimize the score function *S*(Λ), is performed used the augmented Lagrangian algorithm implemented in the nloptr package provided in R (Birgin and Martnez, 2008) using COBYLA (Constrained Optimization BY Linear Approximations) as the local solver (Powell, M. J. D., 1994; Powell, 1998). The method is called successively until the distance between two successive estimates are less than 10^−3^ for all *m* parameters.

The implementation of the CubSFS method also applies to the folded SFS (see the Supplementary Information).

### Estimation of the smoothing parameter and confidence intervals

The smoothness parameter *α* is determined using cross-validation (see Supplementary Figure S4). Briefly, we divide the segregating sites into *K* random groups. For each group we treat the SFS from that specific group as the validation data and the SFS from the remaining groups as the training data. For group *k* we calculate the expected SFS based on the training data E[*ξ*^(*k*)^], and compare to the observed SFS from the validation data, *ξ*^(*k*)^ (*k* = 1, *…, K*). We treat each group as a validation group exactly once and calculate the cross-validation mean square error. We finally estimate the smoothing parameter by minimizing the mean square error

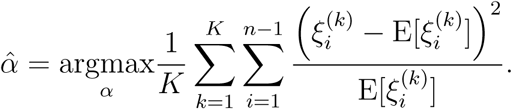

Point-wise confidence intervals can be determined by means of bootstrapping from a multinomial distribution defined from the observed SFS and the number of monomorphic sites. For each bootstrapped SFS we find 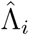, and the lower and upper limits of the confidence interval at time *t* is then determined as the 2.5% and the 97.5% quantile, respectively, of 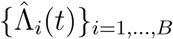, where *B* is the number of bootstrap samples. The median is given by the 50% quantile of the bootstrap samples. We choose *B* = 200.

## Validation and data analysis

Four different models are used to validate the CubSFS method: Exponential growth, two epochs, a bottleneck, and a zigzag model (Schiffels and Durbin, 2014). Recall that we consider 200 sequences of length 5 · 10^8^ and the number of segregating sites is 6 · 10^6^.

In order to apply the CubSFS method we must choose the number and placement of the points *t*_1_, *…, t*_*m*_. We place the points using the same procedure as Li and Durbin (2011; see Supplementary Information eq. (33)). We use a number of points *m* + 1 equal to 4, 7, 10 or 15, and the last time point *t*_*m*_ is placed at 0.5, 1 or 1.5 times the expected time to the most recent common ancestor. For the zigzag model, further results are produced for 15, 20 and 30 time points placed according to knowledge of the model.

The SFS for the 9 different populations used by Liu and Fu (2015) are kindly provided by the authors, and we used the CubSFS method for estimating the changes in populations size back in time. We set the expected time to most recent common ancestor to 600 thousand years ago, and let the last time point be either 1, 1.5 or 2 times 600 thousand years ago. The time points are placed according to equation (33) of the Supplementary Information using a total of 15, 20 or 30 time points. The time points are transformed into coalescent units prior to running the CubSFS method by means of *N*(0) = 10, 000 and a generation time of 24 years, leaving the last time point *t*_*m*_ in coalescent units at 1.25, 1.875(= E[*T*_*MRCA*_]) or 2.5 back in time.

The final results are transformed from coalescent units back to calendar years using a total of *G* = 3, 234.83 Mbp and a mutation probability of *µ* = 1.2 · 10^−8^ per generation per bp (see the Supplementary Information).

### Akaike’s Information Criteria

The final choice of parameter setting is chosen among all suitable parameter settings using the Akaike’s Information Criteria (AIC): The number of free parameters are 2*m* and the likelihood of the observed SFS is assessed through the multinomial distribution defined by the probability of observing a site with *i* derived alleles *p*_*i*_(Λ) (see equation (2)). This probability is either determined from the estimated integrated intensity, or from the points-wise median of the bootstrap samples (see the Supplementary Information)

## Discussion

In this paper we have assumed a Kingmans coalescent in a panmictic population. Under these assumptions the reciprocal coalescent rate is in fact the effective population size. However, a panmictic population is very rare, and whether the change in population size is possible to identify using the SFS is still under debate. Mazet et al. (2016) takes the interpretation of the reciprocal coalescent rate a step further by loosing the panmictic assumption. They find that the estimated coalescent rate can be explained by two different scenarios: Either changes in population size for one panmictic population, or migration between multiple populations with constant population sizes.

The CubSFS method estimates the changes in coalescent rate dependent only on prior defined time points placed back in time on a coalescent scale.

A basic assumption of our roughness penalty approach is that the population size is slowly varying. This assumption is valid for the exponential growth model, and we believe this is the main reason why this model is inferred particularly well. For the discontinuous models, increasing the number of time points seems to enable more specific timing of the instantaneous changes in population size. With only a few, i.e. 4 or 7, time points the dependencies between time points is high. However, with more than 10 or 15 time points the CubSFS method is able to evaluate both instantaneous changes in population size and confidence limits.

The ability to estimate the population size depends on the placement of the time points. Placing the last time point before the expected time to the most recent common ancestor may induce additional fluctuations in the recent past. The time points have to capture the whole period of which the SFS provides information. The CubSFS method provides plausible estimates for well placed time points, even when reaching beyond the time to the most recent common ancestor. In this case the CubSFS method recognises the lack of information after the time to most recent common ancestor by inferring a constant population size.

The score function (equation (1)) is composed of two elements, which both of them may be changed to accomidate any prior knowledge or different scenarios. The first term, measuring the similarities between the observed data and the expected from the model specified by the spline, can include any other summary statistics, i.e. LD patterns or runs of homozygosity. Hence dependencies between sites and quality of the observed data can be included in the estimation similar to the PopSizeABC (Boitard et al., 2016). The main advantage of our approach is that we avoid simulations by detailed analytical considerations.

Future studies would be able to build on the joint SFS of multiple populations, using the covariance between population specific SFSs as a measure of similarity, enabling inference of the different changes in size of the populations. However, adding admixture and migration to this set-up, as well as analysing the analytical expression of the covariance within changing population sizes is a topic for further research.

Likewise, the second term of the score function, i.e. the regularisation term, can be adjusted. Here we use the *L*^2^ norm of the smoothing spline. However, other functions can be used such as the absolute value of the smoothing spline (the *L*^1^ norm). The choice of regularisation will affect the smoothness of the estimated spline.

Green and Silvermann (1994) discuss the fact that using the *L*^2^ norm for regularisation is similar to assigning a Gaussian prior to the space of all smooth function in a Bayesian setting (see Green and Silvermann (1994), section 3.8). Hence, our method is similar to those developed by Palacios et. al. (Palacios and Minin, 2013; Palacios et al., 2015), without the need to infer the gene genealogies.

The last part of the score function is the amount of smoothing, and here we use a cross-validation technique on the segregating sites to estimate the smoothing parameter. Other methods can be used, such as the leave-one-out theory used in kernel estimation. However such techniques may depend on the sample size *n*, of which the cross-validation technique is independent.

## Software Availability

The CubSFS method is implemented in R and is available upon request (please contact BLW at). An R package will be available soon.

## Acknowledgement

We thank X. Liu and Y.-X. Fu for providing the SFS of the 9 populations. We also thank J. Terhorst and Y. Song for providing key insight into the SMC++ method. BLW is funded by The Lundbeck Foundation Initiative for Integrative Psychiatric Research, iPSYCH, Denmark. Grant number R155-2014-1724.

## Author Contribution

AH suggested the roughness penalty approach and developed the proof of Theorem 1. BLW developed the mathematical setting of the cubic spline, implemented the CubSFS method, and conducted the analyses. BLW and AH wrote the paper.

## Competing financial interests

The authors declare no competing financial interests.

